# Pre-Training for Large-Scale Functional Connectome Fingerprinting Supports Generalization and Transfer Learning in Functional Neuroimaging

**DOI:** 10.1101/2024.02.02.578642

**Authors:** Mattson Ogg, Lindsey Kitchell

## Abstract

Functional MRI currently supports a limited application space stemming from modest dataset sizes, large interindividual variability and heterogeneity among scanning protocols. These constraints have made it difficult for fMRI researchers to take advantage of modern deep-learning tools that have revolutionized other fields such as NLP, speech transcription, and image recognition. To address these issues, we scaled up functional connectome fingerprinting as a neural network pre-training task, drawing inspiration from speaker recognition research, to learn a generalizable representation of brain function. This approach sets a new high-water mark for neural fingerprinting on a previously unseen scale, across many popular public fMRI datasets (individual recognition over held out scan sessions: 94% on MPI-Leipzig, 94% on NKI-Rockland, 73% on OASIS-3, and 99% on HCP). Near-ceiling performance is maintained even when the duration of the evaluation scan is truncated to less than two minutes. We show that this representation can also generalize to support accurate neural fingerprinting for completely new datasets and participants not used in training. Finally, we demonstrate that the representation learned by the network encodes features related to individual variability that supports some transfer learning to new tasks. These results open the door for a new generation of clinical applications based on functional imaging data.

**SIGNIFICANCE STATEMENT:** Deep learning models that leverage the increasing scale of available fMRI data could address fundamental generalization challenges. We drew inspiration from other domains that have successfully used AI to address these problems, namely human language technology, to guide our exploration of the potential for this approach in neuroimaging. Our pre-training approach sets a new high-watermark for functional connectome fingerprinting, achieving very high recognition accuracy across different tasks, scanning sessions, and acquisition parameters, even when the duration of a scan is limited to less than two minutes. We showed that we could re-purpose the representation learned by our model to recognize new individuals from new datasets and to predict new participants’ cognitive performance and traits.

## INTRODUCTION

Neuroimaging has revolutionized our understanding of the human brain. Functional MRI in particular has afforded valuable insights into key areas of cognition including memory (Bonnici et al., 2012), executive functioning (Szameitat et al., 2002), language (Hsu et al., 2017; Pereira et al., 2018), object recognition (Carlson et al., 2003; Haxby et al., 2001), and emotion (Padmala & Pessoa, 2008). Functional MRI has also helped us better understand disruptions that occur in neurological (Nathan et al., 2017) and psychiatric diseases (Kaufmann et al., 2017). Despite this progress it has been difficult to find applications for functional neuroimaging beyond research (Ramduny & Kelly, 2025). Generalizing models and algorithms beyond a particular dataset or scanner (Cai et al., 2019; Nakano et al., 2020; Wang et al., 2022) has traditionally proven very difficult (see Dufumier et al., 2022 for relevant discussion). These limitations follow, at least in part, from two critical constraints: 1) non-trivial interindividual variability in brain function (Gratton et al., 2018), and 2) the small sample sizes (i.e., tens of participants) typical of a single investigation, due to the high cost of acquiring fMRI data (Szucs & Ioannidis, 2020). Furthermore, these constraints compound one another: limited sample sizes make it difficult to adequately map out the key dimensions of interindividual variability that exist in the larger population (Marek et al., 2022).

The fields of image recognition, language and speech processing faced similar roadblocks prior to the resurgence of deep learning (LeCun, Bengio, & Hinton, 2015). The combination of deep learning and increasing data volumes allowed researchers to build models that could learn the structure and content of these domains at a much larger scale (see Chen et al., 2022; Sun et al., 2017; Krizhevsky et al., 2012 for examples). Importantly, after training, these models can be repurposed, or fine-tuned, for new tasks (and data) where the features distilled from large-scale training data are brought to bear on subsequent tasks, achieving high performance even when fine-tuning data is limited (a process called transfer learning; Baevski et al., 2020; Devlin et al., 2019; Tan et al., 2018). This approach has revolutionized these research and application spaces and transformed the way image, audio and text data is used. For example, in speaker recognition, discriminative neural-network representations that are first trained on large quantities of data achieve state of the art performance (Nagrani et al., 2017; Snyder et al., 2018). These models can then be repurposed to address new tasks, including classifying clinical states like Alzheimer’s (Pappagari et al., 2020) and Parkinson’s disease (Moro-Velazquez et al., 2020) from speech samples.

Large-scale data collection efforts for MRI and data-sharing consortia (on the scale of thousands of participants) have been launched to study and overcome similar limitations in the context of neuroimaging (Van Essen et al., 2013; Markiewicz et al., 2021). The increasing scale of neural data has supported different sub-fields of “neural fingerprinting” wherein high-level, individualized representations are derived that could characterize the patterns of someone’s neural activity relative to the variability of the larger population (Kumar et al., 2017; 2018; Wachinger et al., 2014; note these are developed using a given dataset’s anonymized subject identifiers and not anyone’s real-world identities). Functional connectome fingerprinting in particular has a long history as a research topic and has proven to be a powerful representation of brain function that quantifies a person’s brain activity by correlating responses from different brain regions over time (Abbas et al., 2020; Amico & Goñi, 2018; Griffa et al., 2022; Sarar et al., 2021; Van De Ville et al., 2021; Wang et al., 2019). Functional connectome fingerprinting has achieved impressive accuracy for recognizing individuals across tasks (Finn et al., 2015) and can be used to predict cognitive performance (Rosenberg et al., 2016; 2020) and traits of individuals (Greene et al., 2018). Connectome fingerprints have also been suggested as a way to support reliable and personalized clinical diagnostics (Ramduny & Kelly, 2025). While impressive, studies so far have relied on a fairly limited set of hundreds of participants often from only a handful of datasets, far removed from the scale of individuals that may be required to accurately predict cognitive function and mental health variables (Marek et al., 2022; but see Spisak et al., 2023). Additionally, there has been very limited study of the generalization of functional connectome fingerprinting to recognizing individuals from new datasets or to perform new tasks, and deep learning is only starting to be applied in this area (Sarar et al., 2021). Other recent work has adapted self-supervised pre-training methods to train “foundation models” for fMRI (Yang et al., 2025; Caro et al., 2024; d’Ascoli et al., 2025) that might serve a similar function as a warm-start for downstream applications where data is limited. These foundation models prove to be powerful tools, but it is currently unclear how they will be ultimately developed, scaled and deployed in the face of heterogeneous fMRI scanning hyperparameters. Specifically, most approaches predict hemodynamic timeseries but differing TRs in downstream datasets could cause misalignment. We take a different approach, relying on the long, productive history of (supervised) connectome fingerprinting as a pre-training task, which may serve as a useful springboard to reliable and personalized clinical applications (Ramduny & Kelly, 2025).

Functional MRI is approaching a point where breakthroughs that combine deep learning and large datasets can be realized thanks to expanding database efforts (Markiewicz et al., 2021) and advances in standardized preprocessing tools (Esteban et al., 2019). To explore this, we pre-trained neural network representations, which we call ‘brain2vec’ for short, that would learn useful features from large, composite datasets for later use on entirely new data and tasks. We based our experiments on the demonstrated utility of functional connectome fingerprinting (Finn et al., 2015) and drew inspiration from work on speaker recognition (Nagrani et al., 2017; Snyder et al., 2018) in developing this approach.

## METHODS

Functional connectome fingerprinting illuminates a core question in cognitive neuroscience: how can we quantify the variability that exists among individuals and their unique cognitive patterns? To meet this challenge, we looked to the task of speaker recognition, which has made progress on related problems in the speech domain, to guide our methodology (primarily Nagrani et al., 2017, but also Snyder et al., 2018 and to a lesser degree Baevski et al., 2020). In speaker recognition, large-scale supervised pre-training has yielded generalizable and transferable representations. There are several similarities that make this comparison particularly useful: First, unique subject identifiers are available in just about every fMRI dataset and these provide a straightforward target for supervised classification during pre-training (similar to speaker recognition). Second, forcing the model to distinguish high numbers of unique participants requires the model to learn nuanced features about individual variability in order to accurately distinguish participants. Therefore, the features that the model learns will likely be important in other datasets (similar to transfer learning of pre-trained speaker recognition embeddings to ‘speaker verification’ tasks for new individuals). Third, fMRI data from participants performing different tasks in the scanner increases pre-training difficulty and encourages generalizability by forcing the model to learn representations of individuals that are independent of a given pattern of task responses (similar to text-independent speaker recognition). Fourth, some robustness to particular scanners or recording conditions can be learned by training on data from a diverse set of scanning facilities (similar to training on data from different audio domains and datasets, e.g., phone conversations, reading, interviews). Fifth, the temporal nature of fMRI is not dissimilar from speech data: both time series can be sub-sampled to generate additional training examples and improve robustness for future samples where duration is limited. Finally, where multiple scanning sessions are present (e.g., from different functional scanning runs but ideally from separate scanning days), data can be completely held out to form a validation set (similar to held out audio recordings).

### Data and preprocessing

We compiled training (HCP1200, Van Essen et al., 2013; NKI-Rockland, Nooner et al., 2012; MPI-Leipzig Mind-Brain-Body, Babayan et al., 2019, Mendes et al., 2019; 1000 Functional Connectomes, Biswal et al., 2010; AOMIC ID1000, Snoek et al., 2021; OASIS-3, LaMontagne et al., 2019) and transfer learning (AOMIC PIOP1 and PIOP2; LA5c, Poldrack et al., 2016; and data from Setton et al., 2023; Spreng et al., 2022; https://openneuro.org number ds003592) corpora from publicly available fMRI databases (https://db.humanconnectome.org for HCP, https://fcon_1000.projects.nitrc.org for NKI-Rockland and 1000 Functional Connectomes, https://central.xnat.org for OASIS-3, and the rest were acquired from https://openneuro.org). We only retained individuals from OASIS-3 who had a Clinical Dementia Rating (CDR) score of 0 for all scans, to maintain a training corpus of only healthy study participants, as all other studies included in the training set targeted healthy, non-clinical populations. We also only retained data from a subset of 21 sites in the 1000 Functional Connectomes study which had complete data (both a functional scan and a structural scan; see Table 1). We ignored any scans with fewer than 100 TRs. All data were minimally preprocessed to arrive at a version of the data in a common volumetric MNI152 space along with a set of confounds for later processing. We carried out minimal preprocessing in-house using fmriprep (version 20.2.6; Esteban et al., 2019) for all datasets, except for HCP, for which we used the preprocessed MNI152 derivatives released by the human connectome project (Glasser et al., 2013), and the AOMIC datasets (for which we used the publicly provided derivatives generated by fmriprep version 1.4.1). All fmriprep runs were executed with the ‘ignore slice timing’ and ‘susceptibility distortion correction’ flags. Text output generated by fmriprep describing these runs for a representative subject from each dataset is available in the supplemental material.

**Table 1.**
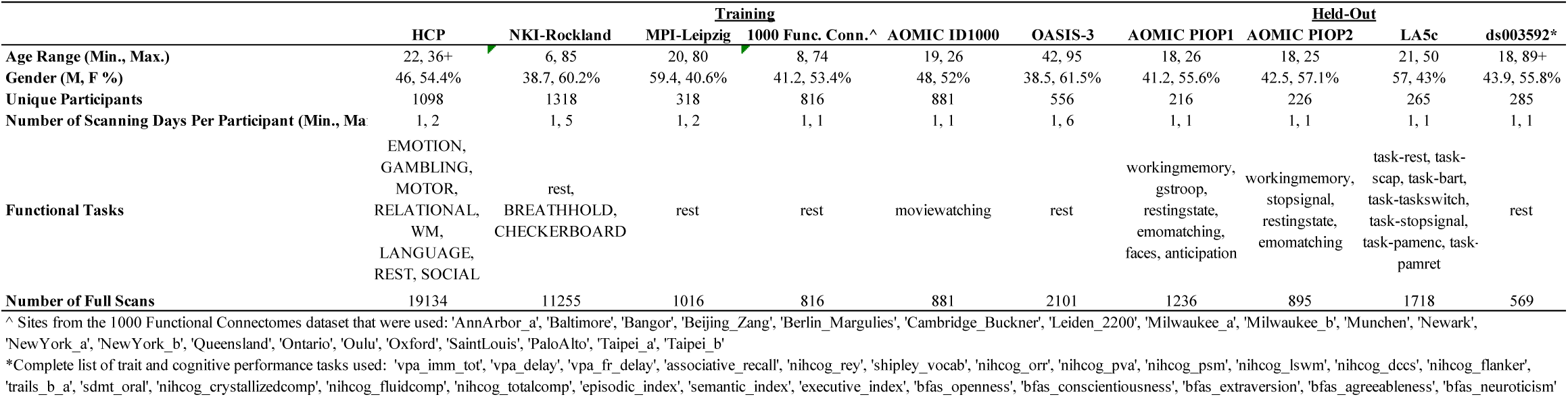
Description of Data.

After minimal preprocessing, all data underwent additional processing via nilearn in python (version 0.8.1). Data were parcellated into a 100-node Schaefer atlas (Schaefer et al., 2018 based on the 17-network atlas, Yeo et al., 2011), standardized (z-scored), detrended and lowpass filtered at 0.08 Hz. We then removed confounds for 12 motion parameters, CSF, white matter and global signal, and then dropped the first 5 samples in the time series. The fMRI time series was then used to derive connectivity matrices (made up of either standard or partial correlation values) for the full scan. For generating additional examples via temporal subsampling, the time series was also subdivided into 100-second segments (with 50% overlap) before deriving additional connectivity matrices. We ignored any data that was not successfully processed by fmriprep or lacked a confounds file. Note that only motion derivatives were available for the HCP data, so preprocessing HCP did not involve CSF or white matter derivatives. A follow-up experiment confirmed that preprocessing the entire training corpus removing only motion derivatives from the time series that were available in HCP (i.e., no CSF or white matter removal) had very limited impact on our results. The control experiment resulted in nearly identical pre-training (validation) performance when all the data were cleaned only with motion parameters. We note that the 100-second temporal subsampling window we used is a very limited duration but aligns with suggested lower-bounds for fMRI analyses that aim to localize activity in time (which is a stricter temporal requirement than ours; Zalesky & Breakspear, 2015; Leonardi & Van De Ville, 2015), and preserves individually identifiable information (Van De Ville et al., 2021).

To monitor training, we constructed a validation set by holding out one scanning session (i.e., one complete day of scanning) from every participant with more than one scan session. These held out scans were separated from the other scans by either days (in the case of HCP) or years (in the case of NKI-Rockland or OASIS-3). The held-out scan was always the last scan for each participant, with the exception of the MPI data. For MPI, the second session, ‘ses-02’ (their Neuroanatomy & Connectivity Protocol), contained significantly more functional data than the first session, so we chose the second session for training and the first session, ‘ses-01’ was used for validation. Note that partitioning the validation data this way means the HCP portion of the validation data involves many tasks not seen during training, since different task were done in the scanner across days. The validation set always involved the full duration scans, ignoring the sub-sampled scans, except for follow up analyses where we investigated model accuracy using the connectivity matrices corresponding to the first 100-second time window. Note that some participants (from NKI-Rockland or OASIS-3) or datasets (1000 Functional Connectomes, AOMIC ID1000) had only one scan. Therefore, these participants only appear in the training dataset and were not used for validation.

### Model architecture and pre-training

We first trained feedforward neural network models to identify individuals in the training dataset from network connectivity matrices derived from their functional MRI scans. We explored two model architectures. The ‘shallow,’ or ‘1 layer’ model architecture comprised a single linear block (made up of a linear layer, leaky-ReLU activation, batch-norm, and dropout) followed by a linear output layer with one activation unit for each participant (4987 units). The ‘deep,’ or ‘4 layer’ model architecture comprised four linear blocks followed by a linear embedding layer (made up of a linear layer, leaky-ReLU activation, and batch-norm) and then a linear output layer (Figure 1). The single linear block in the shallow model doubled as the embedding layer, with the embedding taken before the dropout layer. The input layer of the deep network was four times the size of the embedding layer with the number of activation units in the deep model halved with every subsequent linear block, until the embedding layer in order to encourage the model to distill a more abstract representation at the deeper layers. This also kept the dimensionality of the embeddings from the deep model and the shallow models consistent. All linear layers were initialized with a Kaiming normal function, batch-norm layers had a momentum of 0.1, and did not have learnable affine parameters. Leaky-ReLU layers had operations executed in-place. We used pytorch (version 1.9.1) and pytorch-lightning (version 1.5.10) for neural network training and fine-tuning.

**Figure 1:**
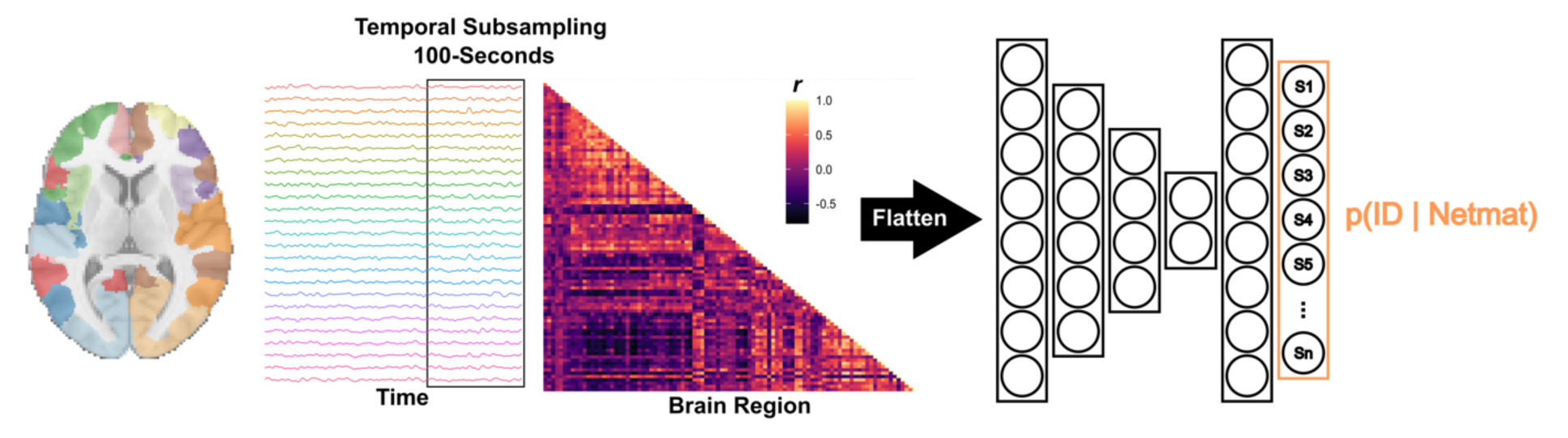
Schematic of the subject classification pipeline. FMRI data are parcellated into a 100-node atlas, and the time series is sub-sampled, converted into a network connectivity matrix and fed into the neural network. Here we show a standard correlation netmat and a deep ‘4 layer’ (plus embedding layer) neural network configuration. The network’s objective is to identify the participant ID with which the network connectivity matrix was associated.

Models were trained for 50 epochs to minimize cross-entropy loss with a learning rate of 0.0001 using an Adam optimizer (weight decay = 0.0001) and a batch size of 64. If there was no improvement in validation loss for three epochs, the learning rate was reduced. Between different numbers of functional sessions and different scan durations, participants had different numbers of training examples. Thus, we applied a class weighting in our loss function to help normalize this. Participants (i.e., the target classes) with fewer training examples received a higher weight, and participants with more training examples had a reduced weight. We evaluated above-chance performance (relative to the number unique individuals in each validation set) using binomial tests, Bonferroni corrected for the number of models we trained (50), and the number of validation tests we ran (5 total: overall performance as well as performance for each of 4 validation datasets).

Training runs encompassed a sweep across different model configurations and hyperparameters. We examined the effect of using different correlations in the input connectivity matrices (standard or partial correlations), model depth (shallow or deep model architectures), model size (embeddings of 1024, 512, 256, 128 units), and dropout (0%, 25% or 50%). Models were compared based on their performance after 50 epochs.

### Ablation analyses

In order to understand which brain regions were most essential for the model’s representation of individual variability, we removed (ablated) portions of the input connectivity matrix and recorded any corresponding changes in validation accuracy. The ablation analysis zero-ed out the rows and columns of the connectivity matrices that corresponded to regions within each of the 17 Yeo networks before presenting them to the model for inference (drawing inspiration from the saliency map analysis run by Lee et al., 2022). For each of the 17 network ablations we recorded validation accuracy and compared this with performance on the un-ablated data to obtain a delta for each ablation. These differences were then normalized between 0 and 1 by dividing by the delta for the network with the highest difference between ablated and un-ablated performance. Note we focus on networks here because the models were fairly robust to zero-ing out individual nodes (brain regions), thus producing very little change in performance.

### Baseline comparison approaches

We used two baseline approaches to understand how our model’s performance fared relative to standard methods (which we describe as the “correlation baseline method” below), and to understand the usefulness of our neural network’s intermediate representation layers (which we describe as the “fully connected baseline method” below). Baseline methods were applied to all the features that were input to our models (100-node input connectivity matrices made up of standard or partial correlations). Our correlation baseline method was an implementation of the method described by Finn and colleagues (2015). This method recognizes individuals by correlating connectivity matrices in the validation set to all other connectivity matrices in the training data. If the highest correlation was with a scan from the same participant this was marked as correct. Our implementation of this approach was carried out within-dataset for computational efficiency (many fewer n-by-n comparisons), but we note this is slightly favorable to this baseline method (eliminates the possibility of cross-corpus false confusions, which is possible for the neural network methods). Correlation methods were performed only on the connectivity matrices from the full-duration scans (i.e., setting aside 100-second sub-sampled examples), with the exception of the analysis where correlation accuracy was assessed using the first 100-second sample of each validation scan.

The fully connected baseline method involved training models that comprised a single fully connected linear layer between the input connectivity matrices and the 4987-unit output layer. Otherwise, these models underwent the same training procedure as the shallow and deep models. Note that at a high level this fully connected baseline approach is similar to connectome predictive modeling (Shen et al., 2017) only we are using a fully connected linear neural network layer to learn the important edges for a given task. A neural network approach here also allows for a matched comparison with our shallow and deep models (and later with models that fine tune brain2vec embeddings), because we can directly match hyperparameters (training data, learning rates and epochs etc.).

### Transfer-learning experiments: Identification of new individuals

We used data from four held out datasets (AOMIC PIOP1, AOMIC PIOP2, LA5c and ds003592) not used during training to understand how generalizable our representations were for distinguishing new participants and data. After preprocessing these fMRI data using the same pipeline above, we sent each (full-scan and sub-sampled) connectivity matrix through our pre-trained models to extract the brain2vec model’s embedding. Note that the models were frozen at this stage and the layers of the model were not updated during this step. We then fine-tuned these embeddings to predict which individual a given training example belonged to, by learning a fully connected linear layer between the model embeddings and a new set of output units, one corresponding to each new participant in these datasets. Importantly, this was essentially a ‘linear-readout’ experiment where we merely updated the mapping between the model embedding and a new set of output units corresponding to these new participants, while the rest of the model was held constant. Thus, successful identification of new subjects had to rely on information contained only within the model embeddings that were pre-trained on the initial training corpus. These identification generalization experiments were carried out for each dataset individually. Participants in these datasets underwent multiple resting state or task scanning runs. We used cross-validation over scanning runs to assess model performance by iteratively holding out each of these runs for testing and training the fully connected output layer on the remaining runs. This fully connected layer was trained for 50 epochs using a cross-entropy training objective, stochastic gradient descent (SGD; momentum = 0.9, weight decay = 0.001) and a learning rate of 0.001. Again, we applied a class (i.e., subject) weighting in the objective to help offset any data imbalance (some participants had missing scans). After 50 epochs for learning the linear mapping between the embedding and the subject classes, the model was tested on how accurately it could recognize the individuals in this dataset based on the held-out scans. We used full-duration and sub-sampled scans as training examples, but only tested on full-duration scans for the held-out runs. We evaluated above-chance performance (relative to the number of unique individuals in each dataset) using binomial tests, Bonferroni corrected for the number of runs in each dataset that we cross validated over and the number of experiments we ran (114; 6 identification methods times 19 held out scans).

Baseline methods were the same as those used for pre-training, only applied to these new datasets. The correlation method matched each held-out scan to the other scans in these corpora by calculating the correlation between the connectivity matrices and taking the subject ID corresponding to the scan with the highest correlation as the model’s prediction. Again, this always involved the full duration scans except when we used the 100-second duration scans as a test of this method under a limited duration constraint. The fully-connected baseline used the same training procedure as above, only here we replaced the model embedding as input to the fully-connected layer with the corresponding connectivity matrices. Thus, this baseline learned a linear layer between the connectivity matrices and a set of output units corresponding to each participant. We trained the fully connected baseline on the same full-duration and sub-sampled training manifest that we used to fine-tune the brain2vec embedding.

### Transfer-learning experiments: Predict traits and behavior

We used data from the same four held out datasets (AOMIC PIOP1, AOMIC PIOP2, LA5c and ds003592) not used during training to understand how generalizable our subject-recognition brain2vec representations were to new tasks. We took the same set of model embeddings (i.e., not updated since the initial pre-training phase) and conducted a new set of experiments where we trained new linear output layers to predict different variables reported in these data. Again, we conducted these within each dataset. For the PIOP datasets we analyzed sex, handedness (categorical variables), Raven’s cognitive test scores, age and NEO personality variables (all continuous variables). Among the LA5c data we analyzed gender, clinical diagnosis (both categorical variables) and age (continuous). Within ds003592 we analyzed each participant’s young or old age grouping, study site, gender (all categorical), as well as age, cognitive (mostly NIH battery), big five personality variables (Big Five Aspects Scale) and MMSE (all continuous, see Table 1), thus a similar set of cognitive dimensions as the PIOP datasets.

These new fully connected layers were trained on data from all participants except one who was held out for testing and we iterated through holding out each participant’s data in turn (i.e., leave-one-out-cross-validation). Models were all trained for 10 epochs with a learning rate of 0.005, using SGD as above. We did not use sub-sampled scans for these experiments because there were many more training examples per output class. Experiments with discrete, categorical variables (e.g., gender, study site, diagnosis) were trained to minimize a cross-entropy loss function (again with class-wise weighting) with one output unit for each of the levels of a given variable (e.g., site 1 or site 2). Experiments with continuous variables (e.g., age, task performance, personality scales) were trained to minimize a mean-squared error loss function with a single output unit. Since we tested on all scans for each participant, we assessed performance for each task for each scanning run. Regression performance was assessed by calculating the correlation between the actual values for held out participants and the model’s predictions (chance level being a correlation of zero, Bonferroni corrected by 990: the number of models assessed times the number of held out scans times the number of tasks we evaluated), while classification performance was assessed via accuracy (chance level relative to the number of classes, Bonferroni corrected the same way for 180). The baseline, fully connected models were trained using the same procedure and again, the only difference was that the training procedure learned a linear layer between the original connectivity matrix and the output.

### Representational similarity analysis (RSA) to understand cognitive dimensions captured by model embeddings

Representational similarity analysis can offer another view into the information encoded in a representation, without explicitly defining a training objective. We ran each resting state scan’s full-duration connectivity matrix from ds003592 through our frozen, pre-trained brain2vec models to obtain their embeddings. We then calculated the cosine distance between the embeddings derived from each subject’s scans to obtain network-dissimilarity matrices. Similar functional-connectome-dissimilarity matrices were derived by calculating the cosine distance between the connectivity matrices that would be used as inputs to our models. Finally, we calculated the difference between each subject’s score on different cognitive and personality measures within the dataset to obtain a set of behavior-dissimilarity matrices. Again, we focused on the cognitive and personality variables reported in ds003592 because there was substantial cognitive testing and metadata and the participant group spanned a wide age range. We calculated the full and partial correlations between the behavior-dissimilarity matrices and the connectome and network-dissimilarity matrices to understand how the differences among individuals on these behavioral measures were captured by different representations of the neural data. These were semi-partial correlations that allowed us to examine the relationship between, for example, a behavioral dissimilarity matrix and a model’s dissimilarity matrix while statistically controlling for the rest of the behavioral variables. These RSA correlations were Bonferroni corrected as a function of the number of network (2) or connectome (2) dissimilarity matrices against the behavioral dissimilarity matrices (27; for 108 tests total). We assessed only session-1 scans, but obtained a similar pattern of results based on the session-2 scans. Finally, we restricted our analyses to participants who did not have any missing data among the behavioral measures we targeted, since this was a prerequisite of the semi-partial correlation function we used.

## RESULTS

### Pre-training neural-networks for functional connectome fingerprinting

We compiled a training dataset from six large fMRI datasets (HCP1200, Van Essen et al., 2013; NKI-Rockland, Nooner et al., 2012; MPI-Leipzig Mind-Brain-Body, Babayan et al., 2019, Mendes et al., 2019; 1000 Functional Connectomes, Biswal et al., 2010; AOMIC ID1000, Snoek et al., 2021; OASIS-3, LaMontagne et al., 2019), which comprised data from 4987 unique participants and 21,453 functional runs. These data were pre-processed to distill a functional connectivity matrix (netmat) based on the 100-node Schaefer atlas that was used as input to the model. Netmats across each full functional run along with netmats from 100-second sub-sampled segments derived from those runs were used as training examples (167,200 total training examples). During training, the model’s objective was to correctly match each functional connectivity matrix with one of the corresponding 4987 participant identifiers in the training dataset. We explored a variety of model parameters including full or partial Pearson correlations for the input netmats, model depth (1 or 4 hidden layers), model size (1024, 512, 256 or 128 neurons in the penultimate hidden, or embedding, layer), and various dropout rates (0, 10, 25, and 50% dropout applied to the hidden layers; see Methods for complete training details). To validate performance during training, we held out data from one day of scans in the cases where a participant came in for scans across multiple days (see Methods for further details on model pre-training). These validation scans were separated from the training scans by either days (HCP) or years (NKI, OASIS-3). We retained the model with the highest performance for both regular and partial correlation netmat inputs for further experiments. We refer to these as the ‘brain2vec’ models.

We were able to achieve high performance on the validation data (over 90% accuracy, all greater than chance; Bonferroni corrected *p* < 0.001) across a wide range of hyperparameters (Figure 2). These models created a rich embedding space that clearly separated individuals (Figure 3). In general, input netmats comprising partial correlations and larger embedding layer sizes improved performance, as did applying dropout to the hidden layers (though high dropout rates hurt performance for larger models). Somewhat surprisingly, a shallow architecture with 1 hidden layer out-performed the deeper architectures with 4 hidden layers. The best performing model achieved 97.64% accuracy on the validation data overall. This model comprised one 1024-unit hidden layer with partial correlations as input and with no dropout applied to the hidden layers during training. Performance was highest for the HCP data (10426 correct out of 10442 scans from 1061 participants; 99.9% accuracy) followed by NKI-Rockland (2465 correct out of 2608 scans from 535 participants; 94.5% accuracy), MPI-Leipzig (103 correct out of 109 scans from 109 participants; 94.5% accuracy) and OASIS-3 (433 correct out of 591 scans from 296 participants; 73.3% accuracy). The top performing model using standard correlation netmats as input comprised one 512-unit hidden layer trained with 50% dropout, and achieved performance that was lower than the best partial correlation model (93.3% validation accuracy overall, 95.0% on HCP, 90.3% on NKI-Rockland, 96.3% on MPI and 75.3% on OASIS-3).

**Figure 2:**
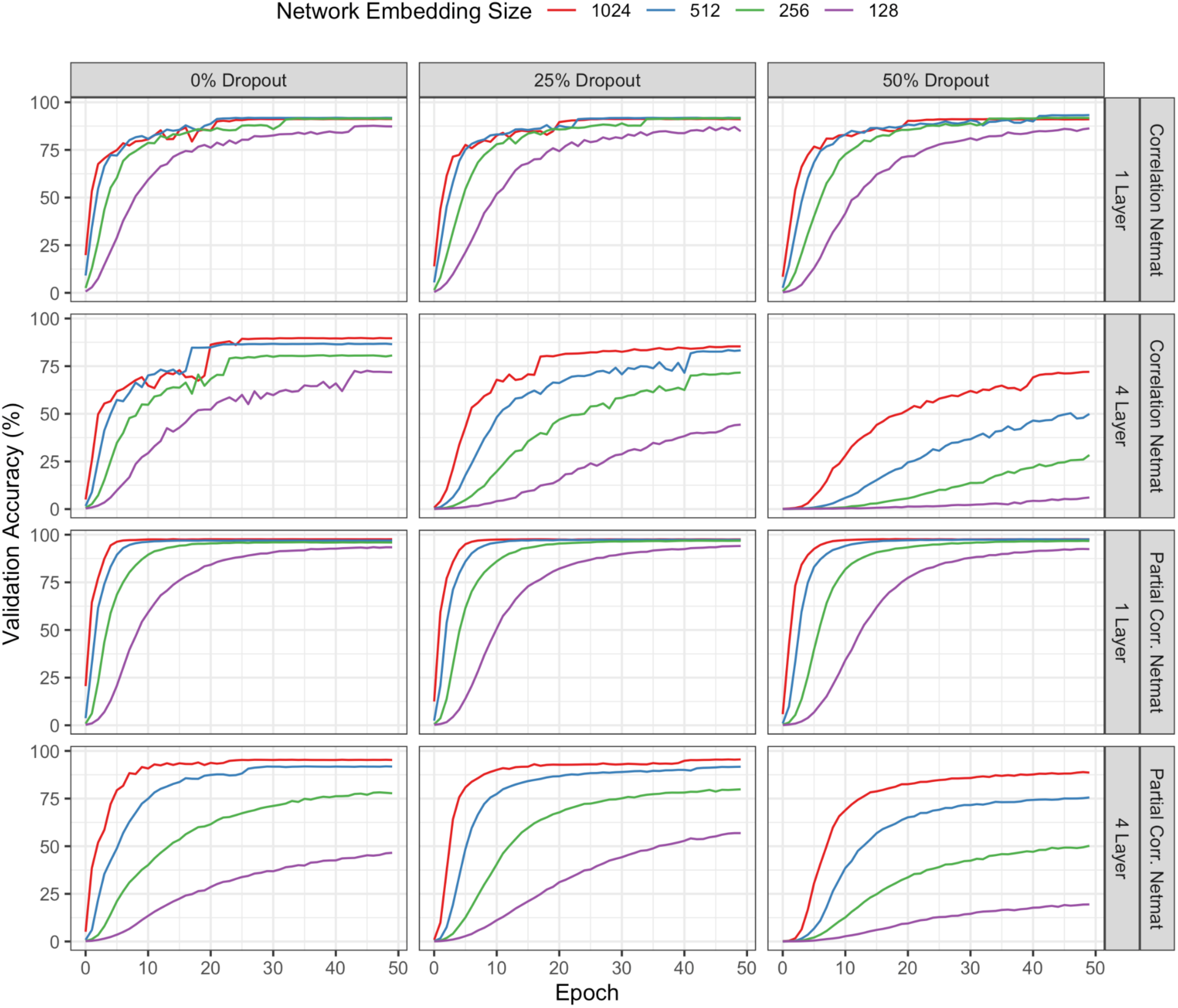
Brain2vec model performance for different hyperparameters assessed on the validation data across training epochs.

**Figure 3:**
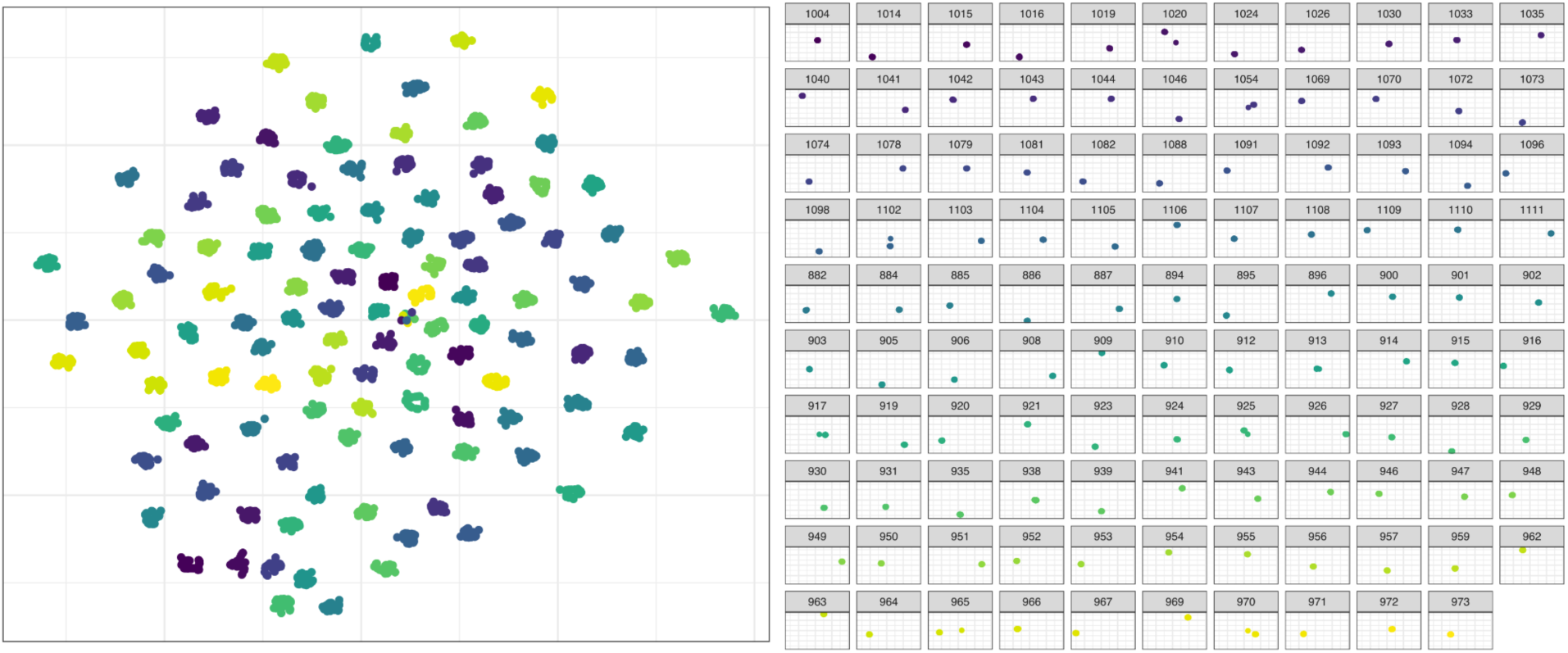
Brain2vec model embeddings for the MPI validation data represented in a two-dimensional space via t-SNE (perplexity = 30, 1000 max iterations). Points are displayed beside one another (left) and faceted by participants (right). All points are colored by which of the 109 participants they belong to (the faceted plot serves as a color key). Points comprise both full duration scans as well as the corresponding 100-second segments. Individuals with multiple clusters or stray points in this subspace indicate scans or time windows that may be more dissimilar from the rest of that participant’s data, and likely contribute to less-than-ceiling classification accuracy.

Performance of the best model (using partial correlation netmat inputs) was near ceiling on the HCP validation data even though the held-out HCP scans comprised many new tasks not seen during training. This kind of generalization should be a difficult task for a subject recognition model since it requires learning features that pertain to individuals despite fMRI data being modulated by performing different, unseen tasks. High performance on these scans suggests the model has learned a high-level, generalizable representation of individuals. This high performance on novel tasks could also be due to the high volume of HCP data used in training (many scans, each fairly long), where the scans used for training also involved diverse tasks. Model performance for the best partial correlation input model was nearly the same, only 0.03% less accurate overall, when evaluation was performed on the first 100 seconds of the validation scans rather than their original full 5 to 15 minute duration (97.61% overall, 99.8% on HCP, 93.8% on NKI-Rockland, 94.5% on MPI and 75.8% on OASIS-3). The top performing full-correlation input model fared worse, almost 5% less accurate overall, when data was limited to 100-seconds (88.9% overall, 90.7% on HCP, 86.7% on NKI-Rockland, 93.6% on MPI and 64.5% on OASIS-3).

### Ablation analysis: Localizing identifiability networks

We wanted to understand which brain regions our model relied on to distinguish individuals. To do this, we carried out an ablation analysis where we re-evaluated the validation data while we iterated through each of the 17 networks (Schaefer et al., 2018; Yeo et al., 2011) in the Schaefer brain atlas, censoring (i.e., zeroing out) each of the networks in the input connectivity matrices in turn. We recorded how much validation accuracy decreased as each network was removed and converted this to a normalized importance score (similar to Lee et al., 2022; see Methods for details). Ablation analysis results are presented in Figure 4. This revealed that removing frontoparietal networks (specifically Yeo-networks 14, 10, 6, and 4) involving the default mode and attention networks incurred the largest reduction in participant recognition accuracy (-0.87%-0.79%, -0.73% and -0.68%, respectively) for the best performing model that relied on partial correlation netmats as input. But we note that overall, our models were fairly robust to censoring individual networks. Accuracies changed only slightly (at most <1%) for each network that was removed (see Byrge et al., 2019 for related discussion). Nonetheless, the importance of frontoparietal networks supports previous fingerprinting results that were able to obtain high accuracy when focusing on these areas (Finn et al., 2015). Overall, a similar pattern was observed implicating the importance of default mode and attention networks for the best standard correlation netmat input model, albeit with larger deltas when each network was censored (most influential Yeo-networks: 13,14, 6 and 11, decreased validation performance -24.2%, -18.7%, - 18.4% and -18.0% respectively).

**Figure 4:**
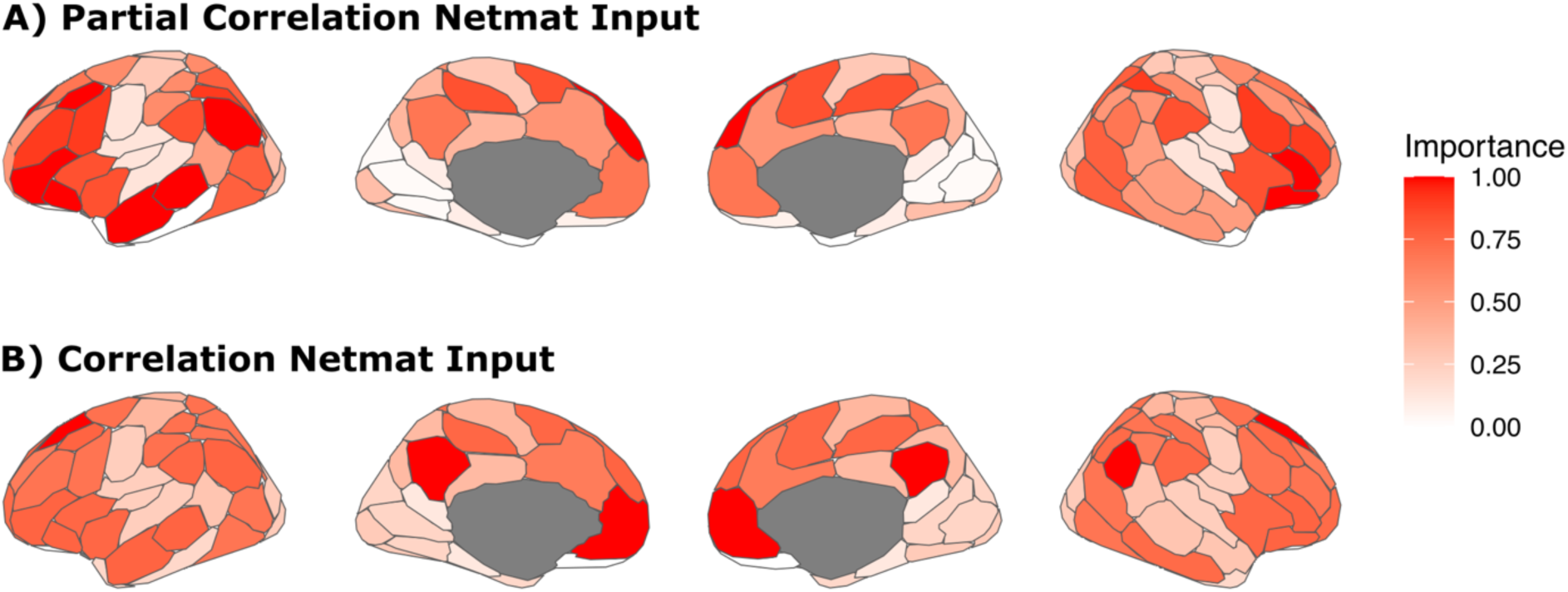
Ablation experiments indicating which networks had the largest impact on validation performance when they were removed for both partial (top) and standard correlation (bottom) netmat inputs. Reduction in validation performance is normalized to derive an overall importance score within each plot (see text for how much validation performance changed for each model).

Neural networks are efficient and powerful learners, and it is difficult to completely rule out whether a model has learned to exploit artifacts in the training data to improve performance. We conducted multiple follow up control experiments to examine the influence of common fMRI artifacts in these data, and found they could not explain our model’s performance. If the models are exploiting motion artifacts unique to individuals, a model trained directly on these artifacts should perform well at distinguishing individuals. However, replacing our fMRI netmat training samples with their corresponding motion confounds (mean and standard deviation of x, y, z and their derivatives over the corresponding time windows) achieved < 4% accuracy on held out days of scans, suggesting that motion is not driving these results. Similarly, if the models are learning to exploit artifacts from anatomical normalization, then spatially filtering the data (a common preprocessing step to mitigate the influence of anatomical artifacts) should reduce performance. Filtering the functional data with an 8mm spatial filter did not result in an appreciable change in validation accuracy for partial correlation netmat input models (97.9% overall and 97.8% on just the first 100-seconds of each scan), but did incur a slight reduction in the performance of standard correlation netmat input models (89.8% overall; 82.9% on just the first 100-seconds of each scan). Overall, these control experiments suggest that artifacts from anatomical normalization are not driving these results.

### Comparison with baseline methods

Finally, our brain2vec model’s performance compares favorably to other methods. For example, correlation-based fingerprinting methods do not perform well on large datasets using partial correlation netmats from the same 100-node parcellation (55.0% overall, 50.6% on HCP, 68.5% on NKI-Rockland, 89.9% on MPI and 66.5% on OASIS-3), but can perform better on smaller datasets (MPI) or those comprised of mostly resting state scans (NKI-Rockland and OASIS-3). The accuracy of the correlation method decreases considerably when using full correlation matrices comprised of 100 nodes (overall validation accuracy: 30.5%). As an additional baseline, we trained a linear read-out layer (i.e., fully connected linear neural network layer between the input and output layer with no hidden nodes) mapping from the input correlation matrices to the 4987 output units (one for each subject identifier) using our full training dataset. This is essentially a neural implementation of connectome predictive modeling, where a fully connected neural network layer is used to learn the most relevant edges in the input connectivity matrices for predicting participant IDs based on the training data. This method also performed well on our validation data, albeit performing below our best performing partial correlation netmat input model (overall validation accuracy: 96.9%; and 74.8% using standard correlation netmat inputs), and worse on the 100-second validation scans (95.6% and 62.9% using partial and standard correlation inputs, respectively). These baseline analyses indicate that the functional connectivity matrices we used as input to our model are very rich features for distinguishing individuals and mapping individual variability. However, neither of these baseline methods produces a generalizable representation that can bring insights that might be learned from the population during pre-training to bear on new tasks.

### Generalizing model representations to identify new subjects from new datasets

An extensible resource in neuroimaging will need to achieve high performance on new datasets containing individuals and tasks not seen during training. Moreover, a useful transfer-learning resource should bring along insights learned from large scale pre-training to aid in analyses of new data. As a first test of how well our functional connectome fingerprinting model’s representations generalize, we conducted a series of transfer learning experiments. We retained the brain2vec models that performed best on the connectome fingerprinting pretraining task relative to the validation data (one based on partial correlation inputs and one based on standard correlation input), froze their weights, and trained a linear read-out layer to classify new individuals based on our model’s frozen representations (see Methods for details). We used data from four new datasets, each containing at least 200 participants: AOMIC PIOP1 and PIOP2, Snoek et al., 2021; LA5c, Poldrack et al., 2016; and data from Setton et al., 2023; Spreng et al., 2022; openneuro.org number ds003592). For each dataset we trained a new fully connected layer to classify these new individuals based on the fingerprinting model’s embeddings. This can be thought of as re-mapping the model’s embeddings to a new set of output units (i.e., one new output unit per participant in the new dataset, replacing the old set of 4987 output units). Each dataset contained multiple scanning runs, often involving different tasks, which we used for cross-validation. A new fully connected output layer was trained (for 50 epochs) using data from all runs except one which we held out for testing, iterating through each task run as the held-out run in turn. We recorded how accurately the model could identify the participants of a given dataset based on each held out run. We call the data used to fine tune the mapping between the model embeddings and the new outputs, “enrollment data,” and the data that is held out to test identification accuracy after fine-tuning the “test data.”

Our model’s representations supported very accurate individual identification on these new datasets (above chance identification on all held out scans, Bonferroni corrected *p* < 0.001; Figure 5). Representations from the best performing partial correlation-input model supported an average of 96.9% identification accuracy for these new participants across all held out scans and datasets (98.5% on PIOP1, 96.2% on PIOP2, 97.6% on LA5c, and 90.8% on ds003592), while representations from the best performing standard correlation-input model supported slightly better (paired-samples Wilcoxon signed rank test: *V* = 17, *p* = 0.003) performance with an average of 98.5% accuracy on these new data (98.7% on PIOP1, 98.4% on PIOP2, 99.5% on LA5c, and 94.9% on ds003592). Datasets with more enrollment data (i.e., a higher number of scans for fine-tuning) achieved better identification performance, with accuracy on PIOP1 ranging from 97% to 100% across held out scans and accuracy on LA5c ranging from 96% to 100%. Performance was slightly lower on ds003592 (90.1% to 95.1%). Low performance on ds003592 could have resulted from limited scanning data or a less diverse set of task runs (all resting state) in the enrollment dataset. In any case, the high performance we observed on the PIOP and LA5c datasets is notable, in that this requires model representations to generalize well across a wide set of tasks, many of which were not seen during training. This high-performance matches or exceeds performance on our original validation data (see previous section) indicating the representation our model learned is generalizable to new participants and scanning protocols.

**Figure 5:**
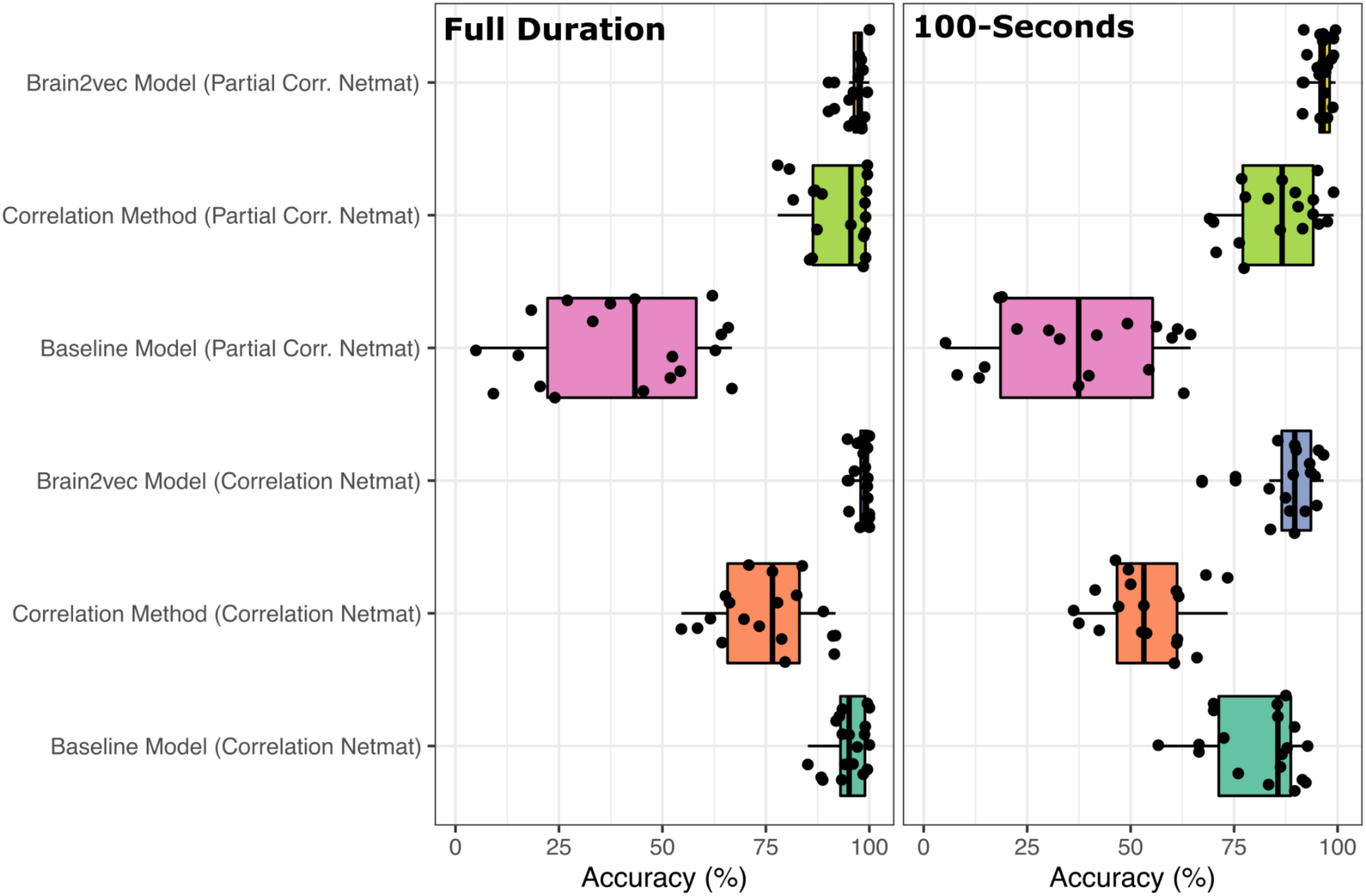
Performance of each identification method on recognizing individuals from new datasets aggregated across all held out scans and datasets. Left: Results based on evaluations of held out scans at their full duration; Right: Results based on evaluations of held out scans when limited to just a 100 second duration.

To understand how beneficial our pre-training strategy was, and whether the intermediate representation of our models encoded useful information for these fine-tuning tasks, we performed two follow up experiments. First, we trained a linear read-out layer to identify participants in each dataset directly from the input netmat correlation matrices using the same set up above (i.e., again holding out each task scanning run in turn). Note that because our model embeddings are much smaller (512 or 1024 dimensions) than these connectivity matrices (4950 dimensions), this allows the fully connected baseline models to have many times more free parameters to perform these tasks. All the fully connected baseline models performed better than chance recognizing individuals from held out scans (Bonferroni-corrected *p* < 0.001). Both of our brain2vec models outperformed their respective fully connected baselines (paired-samples Wilcoxon signed rank test for partial correlation-input model mean = 96.9% vs fully-connected baseline mean = 39.9%: *V* = 190, *p* < 0.001; and for standard correlation-input model mean = 98.5%vs fully-connected baseline mean = 95%: *V* = 168, *p* < 0.001). The partial correlation-input fully connected baseline model performed quite poorly with accuracy ranging from 7% overall on ds003592 to 61.3% overall on LA5c while its counterpart based on standard correlation input netmats performed quite well (88.6% on ds003592 to 97.7% on LA5c). Next, we used the correlation method (described in Finn et al., 2015) to see if we could match each participant’s scan from the held out, test scanning run to another scan from the same participant in the enrollment set based on connectivity matrices (by correlating each test scan’s netmat with all the netmats in the enrollment set and taking the participant identifier belonging to the highest correlated scan as the best match per this method). Both the brain2vec models outperformed these correlation baselines as well, but the difference was only statistically significant for the standard correlation-input model (paired-samples Wilcoxon signed rank test for partial correlation-input model mean = 96.9% vs correlation-method baseline mean = 92%: *V* = 137, *p* = 0.09; and for standard correlation-input model mean = 98.5% vs correlation-method baseline mean = 75%: *V* = 190, *p* < 0.001). The correlation method was able to recognize individuals from all held out scans better than chance (Bonferroni-corrected *p* < 0.001) and, as expected, this method performed well on these smaller datasets because of the limited pools of participants among which to compare correlation matrices (see Ramduny & Kelly, 2025 for discussion). When using partial correlation matrices this method achieved 77.9% to 99.5% across these experiments (using standard correlation matrices as input to this method reduced performance, ranging from 54.6% to 91.8%). Notably, the correlation approach was able to very accurately recognize individuals among the ds003592 scans (98.9% to 99.3% accuracy based on partial correlation netmats) which contained only resting state scans. The correlation method performed worse recognizing individuals for some task runs in datasets with diverse functional runs, however the brain2vec models performed well in these cases.

Finally, this pattern of results was consistent, and the brain2vec models further out performed their respective baselines, when test scans in these held out datasets were limited to a 100-second duration (Figure 5). While all 100-second duration experiments still outperform chance (Bonferroni-corrected *p* < 0.001), performance of all methods based on standard correlation decrease substantially compared to full-duration evaluations. Overall average performance of the brain2vec model based on standard correlation input netmats decreased from 98.4% to 88.2%. Meanwhile the overall average performance of the brain2vec model based on partial correlation input netmats is robust to this truncated scan duration (96.7% to 96.3%). Nonetheless, both brain2vec models outperformed all their respective baselines when duration of the evaluation scans is limited (paired-samples Wilcoxon signed rank tests, all *p* < 0.001) (Figure 5), indicating that pre-training is effective in encoding information from a wider base of participants that improves performance on downstream tasks over an already-powerful netmat representation.

### Generalizing model embeddings to predict traits

Individual variability in functional connectivity patterns have been linked to differences in behavior, traits and cognition (Greene et al., 2018; Rosenberg et al., 2016; Rosenberg et al., 2020). However, inter-individual variability likely comprises a large number of cognitive and neural dimensions and it would be difficult to achieve an adequate sample of the population that can model the relevant brain-behavior dynamics within a single study (Marek et al., 2022). Thus, a powerful application for a generalizable functional fingerprinting resource would be to map the latent space of inter-individual variability within a large sample of the population that encodes relevant features that could be deployed to model brain-behavior associations for new data and applications. If our model has adequately mapped and encoded this latent space, the representations learned by our model could be useful for predicting behavioral performance and individual traits in new datasets.

To test whether our brain2vec fingerprinting model learned representations of individual variability that are useful for predicting subject traits and might improve performance on new tasks, we ran a new set of transfer learning experiments to predict participant traits or performance on several different cognitive tasks, using the same held out datasets (AOMIC PIOP1 and PIOP2, LA5c and ds003592). Again, we froze the network and trained a new fully connected layer to predict traits and behavioral measures based on our model’s embeddings. In these experiments, the output layers were trained to predict subject metadata other than their IDs (see Methods for details). These experiments used a leave-one-subject-out cross-validation scheme. In all cases, the new fully connected output layer was trained for a small number of epochs on data from all subjects except one, and then the model was tested on the held-out participant’s data (see Methods for full description). These experiments involved learning both regression (e.g., for predicting age or performance on a cognitive task) and classification tasks (e.g., for predicting gender or disease status), thus depending on the nature of the data and task the output neurons and training objectives were updated accordingly (to a single neuron trained to optimize mean squared error, or a neuron for each class trained with cross entropy, respectively). Because different tasks performed in the scanner could modulate brain activity in a way that might emphasize useful features of individual variability (Finn et al., 2017; Gratton et al., 2018) or task performance (Greene et al., 2018), we trained the model using all available scan data (here, full-duration scans only) for our subjects. When evaluating the held-out participant’s data we assess model performance for each measure as a function of the in-scanner task the participant performed.

In general, our brain2vec model performance on these non-identification tasks was lower than for recognizing large numbers of individuals (for which our model’s performance was near ceiling), but our models could still predict many task outcome variables well above chance. The best performing pre-trained model that took partial correlation netmat inputs significantly predicted 34 of 40 classification variables (Figure 6) above chance and 22 of 131 regression variables (Figure 7) above chance (all Bonferroni-corrected *p* < 0.05). Its counterpart that took correlation netmat inputs significantly predicted 31 of 40 classification variables above chance and 25 of 131 regression variables above chance (all Bonferroni-corrected *p* < 0.05). Our models performed best on classification tasks. Neither model could predict continuous variables in the PIOP1 and PIOP2 datasets (though classification performance on those datasets was similar to other datasets). Models performed best on the ds003592 data, accurately predicting study-site and age-group (better than 70% accuracy) as well as age (*r* > 0.74, and < 12.3 MAE). We were also able to significantly predict participant performance on different memory and cognitive variables (albeit to a lower degree of accuracy) including performance on the Verbal Paired Association test (delay; *r*=0.28 to 0.42), Associative Recall Paradigm (*r*=0.38 to 0.46), Symbol Digits Modality Task (*r*=0.27 to 0.33), and Trails Making Task (*r*=0.34 to 0.41; all Bonferroni-corrected *p* < 0.05).

**Figure 6:**
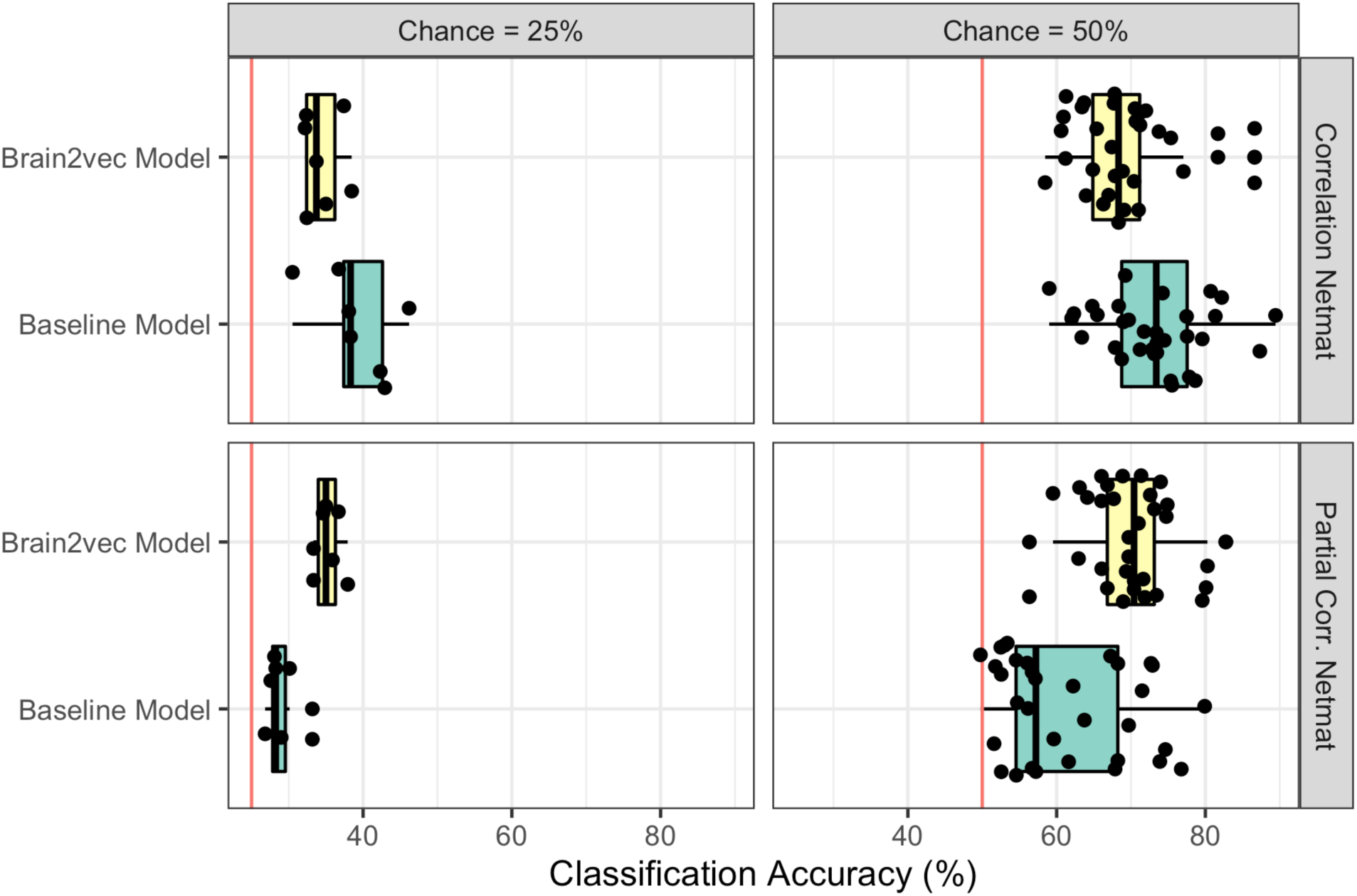
Performance of each identification method predicting categorical variables in new datasets aggregated across all held out scans and datasets, displayed by chance-level. Red lines indicate chance performance.

**Figure 7:**
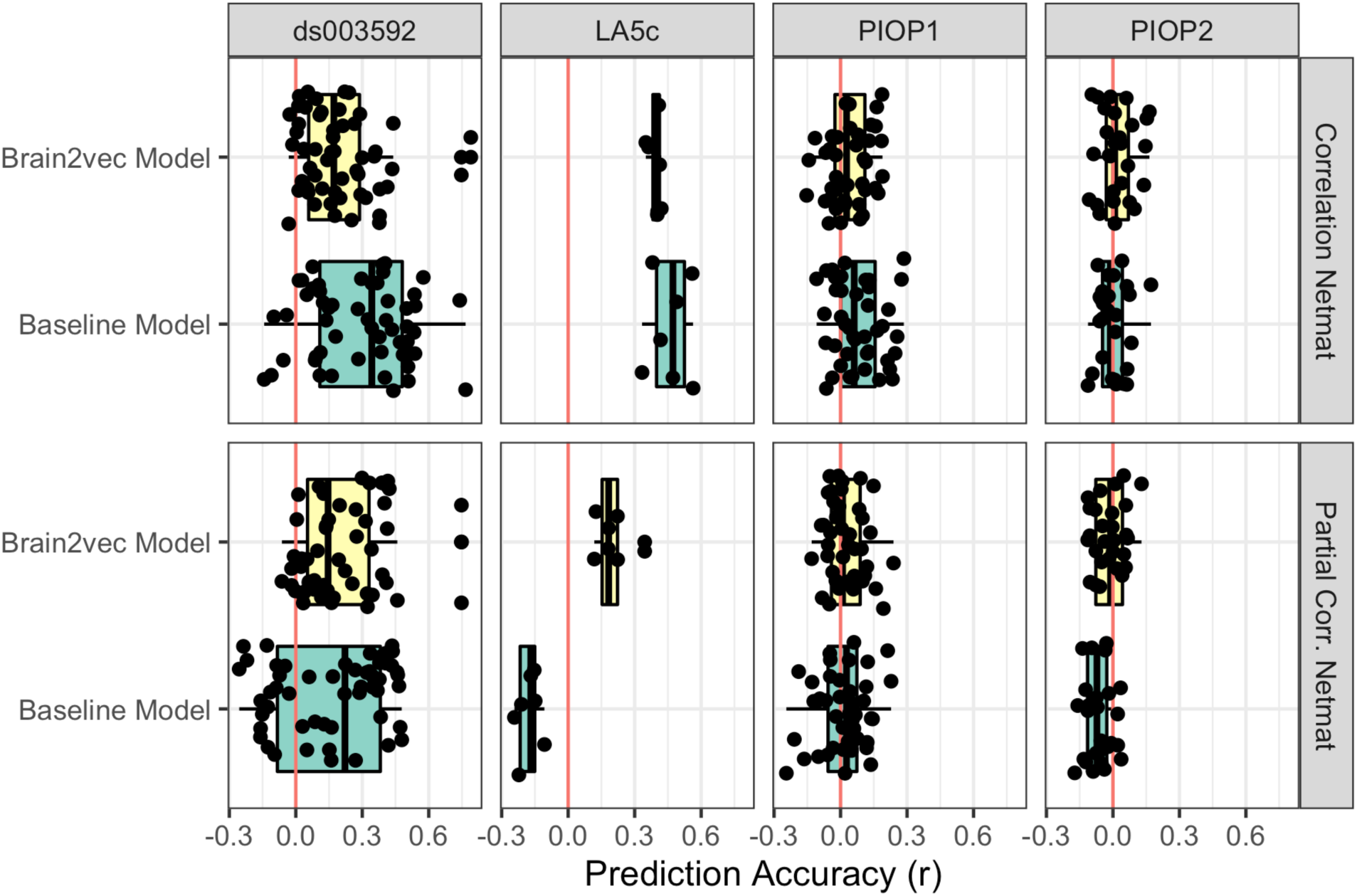
Performance of each identification method predicting continuous variables in new datasets aggregated across all held out scans, displayed by dataset. Red lines indicate chance performance (*r*=0 no correlation between model predictions and true values).

We wanted to determine if our embeddings and pre-training procedure distilled insights from the larger population they were trained on about traits and cognitive performance beyond what could be gleaned from the (already quite powerful) connectivity matrices we used as input. To do this, we trained a series of baseline, fully connected models for a comparison, similar to the generalization experiments recognizing new individuals in new datasets. These models underwent the same procedure, training a single fully connected layer from the flattened connectivity matrices instead of using our model’s frozen, pre-trained embedding as input. Again, note that our model embeddings have a much smaller dimensionality (512 or 1024 dimensions) than the full connectivity matrices (4950 dimensions), and thus have many fewer free parameters for these fine-tuning experiments. Overall, there was only a slight numerical increase in performance from our models vs the baseline models for the categorical, classification tasks, and this difference was not statistically significant (paired-samples Wilcoxon signed rank test: *V* = 1179.5, *p* = 0.05) and no significant difference in performance for the continuous, regression tasks (paired-samples Wilcoxon signed rank test: *V* = 18629, *p* = 0.25). This was because the results for each input feature space were mixed. Our pre-trained model that took partial correlation netmats as input outperformed its fully connected netmat baseline model (paired-samples Wilcoxon signed rank test for classification tasks: *V* = 35, *p* < 0.001; and for regression tasks: *V* = 3309, *p* = 0.02), but the pre-trained model that took Pearson correlation netmat inputs performed significantly *worse* than its baseline (paired-samples Wilcoxon signed rank test for classification tasks: *V* = 658, *p* < 0.001; and for regression tasks: *V* = 6171, *p* < 0.001)

### Representational similarity analysis to understand cognitive dimensions captured by model embeddings

Probe tasks are a useful way to understand the information encoded in a model representation (e.g., Raj et al., 2019; Peri et al., 2020) and demonstrate potential utility for new applications, but these involve many choices including hyperparameters, training targets and optimizations. Representational similarity analysis can provide another view into the information encoded by a network embedding that is less biased by optimization goals and tuning parameters (Mehrer et al., 2020; Kriegeskorte & Kievit, 2013; Ogg & Skerrit-Davis, 2021; Ogg et al., 2024). We used the extensive cognitive and behavioral measures obtained by Setton, Spreng and colleagues (2022; 2022) in ds003592 to carry out a representational similarity analysis that would examine what participant traits or cognitive features our network encoded.

Correlations among the network and functional-connectome-dissimilarity matrices showed only a weak to moderate association with one another. The partial and standard correlation brain2vec model embedding representations were moderately correlated (*r* = 0.35) with one another, and these models were moderately correlated with their respective input feature spaces (*r*=0.52 and *r*=0.33 between netmats and model embeddings that took partial and standard correlation netmats, respectively). The partial and full correlation netmat representations were also moderately correlated (*r*=0.35). This suggests that the pre-training process produces a latent space that differs from the original input space and that different netmat features encode information differently from one another, which could be useful in differential utility for downstream tasks.

Finally, we wanted to understand how the latent space of our models and the input network connectivity matrices encoded traits and cognitive features of individuals. These results are presented in Figure 8. The strongest associations across the neural and behavioral variables were for Age, the Trails Making Task, Verbal Paired Associates and the Flanker task. Most of these associations were common across both the model and netmat features, but the amount by which these representational spaces correlated with behavioral features differed. Notably, many associations between behavioral and neural representations persisted even after statistically controlling for age, which likely introduced some bias related to the cognitive and memory measures in our fine-tuning experiments.

**Figure 8:**
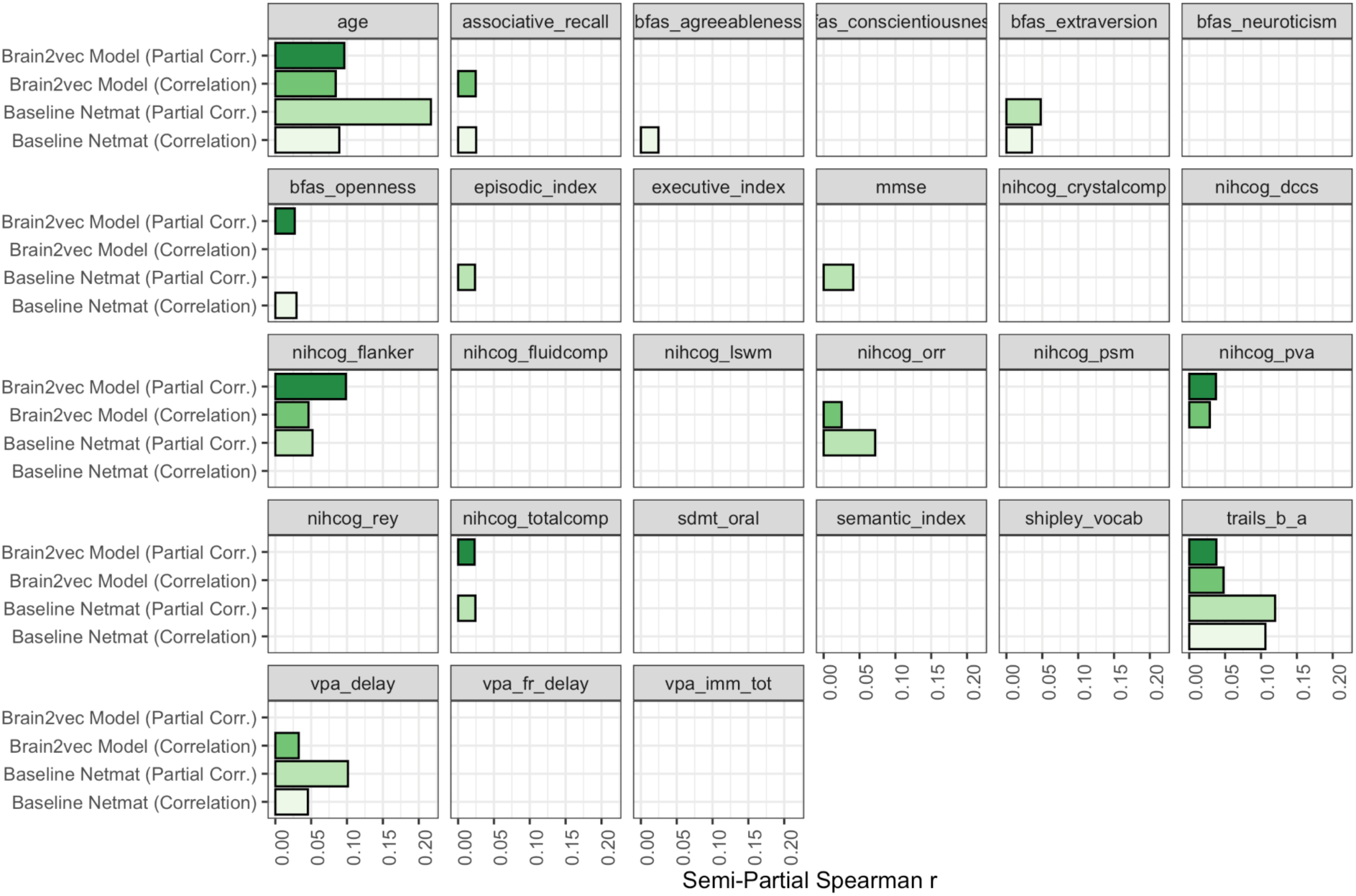
Significant semi-partial correlations between behavioral representations and neural representations after holding all other features constant.

## DISCUSSION

Neuroimaging data has increased in scale, inviting important questions regarding whether these data can be harnessed to distill generalizable representations that could be used for downstream clinical or psychological applications or to gain new research insights. However, this may require a very large number of participants (Button et al., 2013; Helwegen et al., 2023). Here, we sought to pre-train model of neural data that could provide a representation of brain function that is aware of the high dimensional space of individual variability, so that downstream or more data limited applications could perform better with respect to these challenges. Similar insights have been gleaned in other domains like text (Devlin et al., 2019), speech (Amodei et al., 2016) and vision (Krizhevsky et al., 2012) by combining large datasets and pre-training neural network models.

This approach has also been successful in other areas of neuroscience (e.g., Leonardsen et al., 2022; Coon & Ogg, 2025), but generalizable deep learning applications for fMRI have been less well studied, likely due to issues stemming from the diversity of scanning protocols or the high cost of acquiring MRI (Szucs & Ioannidis, 2020).

Our work constitutes a key step towards generalizable representations for fMRI. Where many fMRI applications may be challenged by the large individual variability that exists among population samples (Gratton et al., 2018; Seghier & Price, 2018), we sought to leverage this variability as a resource for generalization. Similarly, we hope our large-scale, pre-trained representation could distill information that could be useful for smaller datasets where it may be difficult or impossible (for example in the case of rare diseases) to get the large sample sizes that may be needed for brain-wide association studies (Marek et al., 2022). The results presented here make some progress on these fronts.

We trained neural networks to recognize a high number of individuals from functional connectivity matrices, achieving high performance among held out scans from known participants. Perhaps more importantly, the model’s representation could be used to classify entirely new participants from datasets not used during pre-training. Finally, this model supported above-chance prediction on new tasks beyond distinguishing individuals, albeit to a lesser accuracy. However, these initial results are encouraging and may invite further exploration in pre-training and fine-tuning models across datasets in fMRI.

Functional connectome fingerprinting has been an active area of research (Abbas et al., 2020; Amico & Goñi, 2018; Finn et al., 2015; Griffa et al., 2022), but these studies have primarily involved modest numbers of individuals from single high-quality datasets. Neural network applications in this area are also not new (Misra et al., 2021; Sarar et al., 2021; Wang et al., 2019), but again, these studies have not looked at repurposing the intermediate representation of these models for new data. We achieve near ceiling accuracy on multiple datasets, even when scan duration is severely limited. However, we note that accuracy among older populations (e.g., the OASIS-3 data) leaves room for improvement. Since we were investigating a large-scale problem we used all available data, regardless of scanner task, but it will be interesting to see if certain tasks or procedures can emphasize specific or different dimensions of individual variability during pre-training or for improving fine-tuning (see Finn et al., 2017 for discussion), or if automated quality control approaches (which is itself an active area of research, see Taylor et al., 2023) to triage data before pre-training further improves performance or generalization. We broaden the scope of this work by looking at multiple datasets together, and creating transferable representations that can encode individual variability to support recognition on new datasets. One interesting observation was that the correlation method we used as a baseline does not scale well to large datasets with thousands of individuals and diverse scanning runs. However, matching scans from individuals across sessions by taking the highest correlation is effective when data is limited to a few hundred participants and this method is especially effective when the task is a resting state scan in both the enrollment and test dataset, as has also been pointed out using a subset of the HCP data (Finn et al., 2015). Here, we demonstrate this on a large scale across more datasets (compare correlation method performance for ds003592, 98.9% to 99.3%, to performance on LA5c, 77.9% to 87.4%). We observed the reverse pattern when training a linear read-out layer to classify participants directly from correlation matrices with no hidden layers: this method performed very well when data is plentiful (e.g., among the large set of training participants, 96.9% validation accuracy in pretraining) but this approach can fail when data is limited (e.g., when datasets were limited to each of the held-out corpora, 7% to 61.3% on the identification transfer learning experiments). Our method of training a generalizable representation performs well both when data is abundant and when it is limited, and also supports accurate individual identification when generalizing across datasets. Finally, our functional connectome fingerprinting models rely on frontoparietal networks (e.g., the default mode network) and implicated a similar set of regions for individual variability as in previous work (Finn et al., 2015).

Our primary goal in this work was to try and develop a transferable representation for different tasks. We take similar cues from where the field is moving, including the prediction of new states (Finn & Rosenberg, 2021; Finn & Constable 2016; Poldrack et al., 2020; Woo et al., 2017). We partially succeed on this front. The representation learned by our model was most useful for supporting tasks similar to those used for pre-training (distinguishing individuals within new datasets), but was not as successful in other non-subject-recognition tasks. For example, our models could be fine-tuned to significantly predict one of four classes of psychiatric disease within the LA5c dataset (32-38% accuracy), but this is far from ceiling, and was not as accurate in some cases as a baseline model (as high as 46% accurate). We note that this reduced performance may at least partially stem from a domain shift. Since the model was pre-trained on a large quantity of healthy individuals, connectivity patterns associated with different neuropsychiatric conditions likely posed an out-of-domain problem for our model. Nonetheless, our model could also be used to significantly predict some cognitive and memory performance variables (e.g., VPA, Associative Recall, Trail Making within ds003592, but not within the PIOP data), reasonably accurately (*r* = 0.28 to 0.45), in the range of some past efforts (Greene et al., 2018) but performing lower than others (Rosenberg et al., 2016; Finn et al., 2015). Nonetheless, our results suggest that for some tasks, even transformations learned via these shallow neural networks can learn a compact representation of a high dimensional latent space that is generalizable to new tasks and data. In fact, our shallow network that performed well in this work is similar to early NLP architectures (e.g., word2vec, Mikolov et al., 2013). In our fine-tuning experiments, this means our model representations (comprising 512 or 1024 dimensions) often performed as well as or better than the baseline models (with 4950 dimensions) using far fewer free parameters.

Age is encoded in our model’s representations and has a large influence on the downstream tasks, but does not explain all the variance captured by our model. For example, AOMIC PIOP1 and PIOP2 have limited age ranges and our model could support accurate recognition of these individuals and other categorical demographic features. Our model also performed well among the HCP participants, who are all young adults and fall within a limited age range. Our representational similarity analysis also highlights that our model encodes features related to executive functioning and memory, even when partialing out variance related to age. Still, age was well predicted by fine tuning our model, and accuracy on downstream tasks that were not related to subject identification was lower, emphasizing more work is needed here. Future work should better understand how features are encoded in these neural network layers and how to re-weight features learned in an initial training round. Indeed, since shallow architectures performed best in pre-training most of our experiments focused on these, but an interesting approach would be to only freeze or fine tune some layers of deeper models.

We used as input to our model an already powerful network connectivity matrix representation that summarizes a window of a time series allowing for more harmonization across disparate scanning protocols. This intermediate representation made it possible to homogenize (at least to some degree) the data from different studies so they could be compiled for a large training run. Indeed, this offered new insights: the network likely learned characteristics of these different site or dataset biases, as evidenced by the success of fine-tuning our neural network representation to discern from which site a participant was scanned among ds003592.

Finally, we chose functional connectome fingerprinting as a supervised pretraining task, because of its demonstrated utility in different areas of neuroscience (Greene et al., 2018), and because a similar approach has some clinical utility using speech (Pappagari et al., 2020; Moro-Velazquez et al., 2020). However, this is not necessarily an optimal pre-training objective, and different downstream tasks may require different pre-training methods. Brain-age prediction, for example, is another supervised task using labels present in most public data that may distill a representation of brain function that is clinically useful (see Leonardsen et al., 2022). Unsupervised methods have also proven to be powerful for leveraging even more pretraining data in text (Devlin et al., 2019) and speech (Baevski et al., 2020) processing tasks. These methods could be very powerful if applied to neuroimaging data.

## CONCLUSION

This work pre-trained a flexible generalizable representation for fMRI data. Using connectivity matrices as input allowed us to aggregate over many scanning protocols, and to scale up a critical neuroscience task (connectome fingerprinting) to support a new data-driven neural-network resource. Along the way we set a high-water mark for connectome fingerprinting. We hope these results stimulate the field to think of how to flexibly repurpose machine learning representations for new data and tasks, and stimulate new research on data and neural-network driven biomarkers for neuroscience.

## ACKNOWLEDGEMENTS

The authors acknowledge the support from the Independent Research and Development (IRAD) Fund from the Research and Exploratory Development Mission Area and the National Health Mission Area of the Johns Hopkins University Applied Physics Laboratory. We thank Jordan Matelsky for providing thoughtful comments on this manuscript and for discussions of this work.

This work would not have been possible without the numerous researchers who shared their data publicly. Data were provided in part by the Human Connectome Project, WU-Minn Consortium (Principal Investigators: David Van Essen and Kamil Ugurbil; 1U54MH091657) funded by the 16 NIH Institutes and Centers that support the NIH Blueprint for Neuroscience Research; and by the McDonnell Center for Systems Neuroscience at Washington University. Data were provided in part by OASIS-3: Longitudinal Multimodal Neuroimaging: Principal Investigators: T. Benzinger, D. Marcus, J. Morris; NIH P30 AG066444, P50 AG00561, P30 NS09857781, P01 AG026276, P01 AG003991, R01 AG043434, UL1 TR000448, R01 EB009352. AV-45 doses were provided by Avid Radiopharmaceuticals, a wholly owned subsidiary of Eli Lilly. The MPI-Leipzig Mind-Brain-Body dataset data was obtained from the OpenfMRI database. Its accession number is ds000221. LA5c data was obtained from the OpenfMRI database (https://openfmri.org/dataset/ds000030/). Its accession number is ds000030.

